# Hippocampal-entorhinal transformations in abstract frames of reference

**DOI:** 10.1101/414524

**Authors:** Raphael Kaplan, Karl J Friston

**Affiliations:** Wellcome Centre for Human Neuroimaging, University College London, United Kingdom; Egil and Pauline Braathen and Fred Kavli Centre for Cortical Microcircuits, Kavli Institute for Systems Neuroscience, Norwegian University of Science and Technology, Trondheim, Norway

## Abstract

Knowing how another’s preferences relate to our own is a central aspect of everyday decision-making, yet how the brain performs this transformation is unclear. Here, we ask whether the putative role of the hippocampal-entorhinal system in transforming relative and absolute spatial coordinates during navigation extends to transformations in abstract decision spaces. During fMRI scanning, subjects learned a stranger’s preference for an everyday activity – relative to one of three personally known individuals – and subsequently decided how the stranger’s preference relates to the other two individuals’ preferences. We found that entorhinal cortex/subiculum signals exhibited reference frame-sensitive responses to the absolute distance between the ratings of the stranger and the familiar choice options. In contrast, striatal signals increased when accurately determining the ordinal position of choice options in relation to the stranger. Paralleling its role in navigation, these data implicate the entorhinal/subicular region in assimilating relatively coded knowledge within abstract metric spaces.

## Introduction

Learning other people’s attributes is facilitated by expressing personal preferences *ordinally*– whether we prefer one thing to another–and *metrically*–how much more we prefer one thing over another. Ordinal and metric coding are particularly important when acquiring knowledge about new people. This type of learning involves relating a new person’s attributes to prior beliefs about other people; either by adopting a relative or absolute frame of reference. For instance, imagine you are preparing dinner for a foreign visitor that says, “I like spicy food”. If they are Vietnamese, their preference for spicy food is probably greater than Germanic tastes, despite everyone declaring the same preference.

On one hand, progress has been made in linking the hippocampus to the maintenance of an ordinal sequence or ‘hierarchy’ of personal attributes (Eichenbaum, 2015; Schiller et al., 2015; Kumaran et al., 2012, 2016) and, similarly, category learning (Zeithamova et al., 2008; Mack et al, 2017, 2018). Yet, the neural representation of metrically coded knowledge remains elusive, even though metric coding affords the transformation of knowledge learned via relative and absolute frames of reference.

Clues about the neural computations underlying the transformation of abstract knowledge among frames of reference may come from research on the role of the hippocampal formation in path integration: the process of calculating one’s position by estimating the direction and distance one has travelled from a known point. During path integration, specific sub-regions of the hippocampal formation have been implicated in integrating relative and absolute spatial coordinates during navigation, in order to reach a desired location (McNaughton et al., 2006). In particular, grid cells in entorhinal/subicular areas are selectively active at multiple spatial scales when an animal encounters periodic triangular locations covering the entire environment (Hafting et al., 2005; Boccara et al., 2010), while hippocampal place cells code specific locations in an environment (O’Keefe & Dostrovsky, 1971). Working together with boundary vector cells in entorhinal/subicular areas – that code an environmental boundary at a particular direction and distance (O’Keefe & Burgess, 1996; Hartley et al., 2000; Burgess et al., 2000) – and head direction cells (Taube et al., 1990), spatially-modulated neurons in the hippocampal formation are thought to serve collectively as a cognitive map of the environment (O’Keefe & Nadel, 1978; McNaughton et al., 2006; Hartley et al., 2013).

Notably, recent findings have extended the idea of map-like coding in the entorhinal cortex and subiculum to humans. Human entorhinal/subicular regions respond to both the distance of goal locations (Howard et al., 2014; Chadwick et al., 2015) and discrete abstract relations (Garvert et al., 2017), suggesting that entorhinal/subicular areas might represent abstract knowledge metrically along multiple dimensions. Taken together, these results imply that neural computations in the hippocampal formation related to spatial exploration, may also underlie the integration of information learned in different reference frames during more abstract types of memory-guided decision-making (Kaplan et al., 2017a).

We investigated whether specific brain regions, including sub-regions of the hippocampal formation – like the entorhinal/subicular area – facilitate switching between relative and absolute reference frames during memory-guided decisions. To test this hypothesis, we developed a novel experimental task, where healthy volunteers were first asked to rate 1-9, on a 0-10 scale, how likely (likelihood) they (*self*), a close friend (*friend*), and a typical person (*canonical*) were to partake in a variety of everyday scenarios (e.g., eat spicy food, read a book, cycle to work; Fig. 1A). To allow for strangers with more extreme ratings than the familiar individuals in the fMRI paradigm, subjects were restricted to rating between 1-9. Subsequently, subjects performed a forced-choice fMRI task (Fig. 1B), where they judged the proximity of a *stranger’s* rating relative to the ratings of the self, friend, and canonical individuals, for a given everyday scenario (Fig. 1B). Specifically, on each (self-paced) trial, subjects were shown the preference of a *stranger* on a 0-10 scale, relative to a known person (*anchor*) and subjects had to select which of the two remaining (*non-anchor*) familiar individuals were closest to the *stranger*. After making a decision, a jittered (mean=2.43s) intertrial interval (ITI) period started, where a white fixation point overlaid on a black background appeared on the screen. In our task, it was crucial that the *anchor* was always placed in the middle of the scale, to ensure that subjects used their prior knowledge in order to infer the *stranger’s* absolute preference and form an appropriate mental number line of personal preferences (i.e., remembering the absolute positions of the different preferences on the number line, instead of the visible relative position of the anchor in the middle of the screen with a stranger presented somewhere between 0-10; see Fig. 1C-D). In other words, subjects had to infer the *stranger’s* position in relation to the *anchor’s* ‘true’ rating for that scenario (i.e., the rating they gave for that anchor and scenario during the training period) and mentally rescale this information relative to the closest boundary (0 or 10). Note that this is a non-trivial task because the preferences of *strangers* were not conserved over attributes.

**Figure 1.**
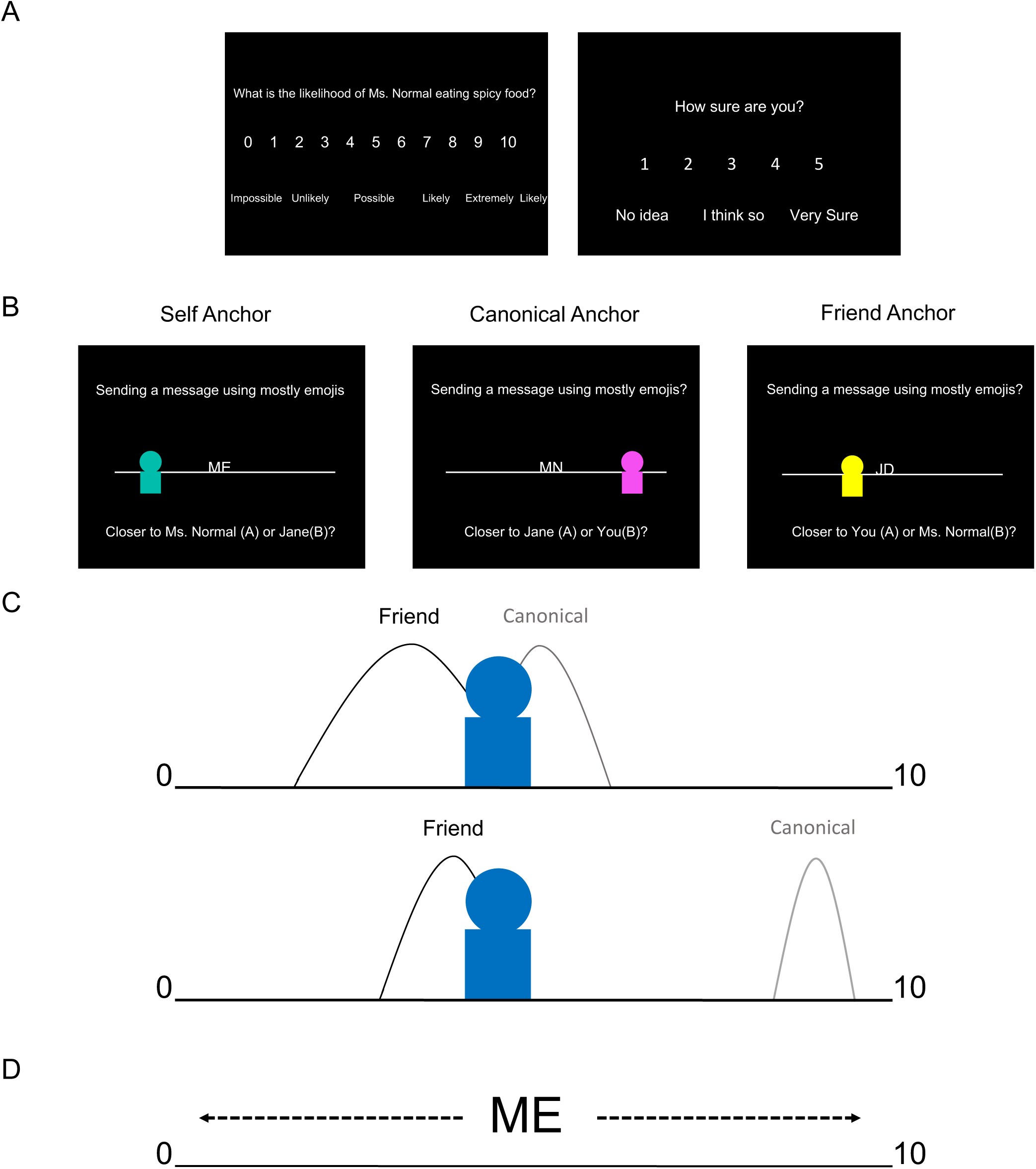
Experiment. *A.* Right before fMRI scanning, subjects were instructed to choose a friend with a different personality of the same gender. Subsequently, subjects rated from 1-9, on a 0-10 scale, how likely (likelihood) they (*self*), a close friend (*friend*), and the typical person (*canonical*) were to partake in a variety of everyday scenarios (e.g., eat spicy food, read a book, cycle to work). To allow for strangers with more extreme ratings than the familiar individuals in the fMRI paradigm, subjects were restricted to rating between 1-9. Subjects also reported their confidence, on a 1-5 scale, for each scenario. *B.* fMRI paradigm. During a forced-choice task, subjects made a decision on the relative proximity of a *stranger*’s likelihood rating for an everyday scenario relative to the likelihood ratings for the *self, friend*, and *canonical* individuals for that same scenario. On each self-paced trial (max. allowed response time 9s), subjects viewed a personal preference for a new *stranger* presented relative to one of the known individuals’ initials (*anchor*) on a number line. Subjects had to determine which one of the two remaining (i.e., non-anchor) familiar individuals was closer to the *stranger’s* rating. Crucially, the anchor individual (e.g., ME=self; MN=canonical ‘Mr./Ms. Normal’; JD=friend’s initial) was always placed in the middle of the scale, ensuring that subjects had to use memory of their ratings made before fMRI scanning to infer the *stranger’s* absolute preference relative to the *anchor’s* true rating and the ends of the number line. Additionally, subjects were instructed that the number line ranged from 0 and 10. For example, if the stranger was ¾ to the right of the anchor with a rating of 9, the participant would infer that the stranger’s rating was ¾ of the way between 9 and 10 (the right boundary of the scale). Consequently, the participant would indicate that the stranger’s rating would be approximately near 10. Notably, the numbers on the scale were not visible during the task. After making a decision, an intertrial interval (mean=2.43s) screen with a white fixation point in the center of a black screen appeared. *C.* Illustration of the behavioral model. Top illustration shows an ambiguous, less discriminable choice, while the bottom illustration shows a straightforward, highly discriminable choice. We quantified the difficulty of discriminating a particular choice by fitting a formal signal detection model, based on the absolute distance between the two choice individuals on the scale and how confident subjects were in their ratings (e.g., comparing the *stranger* rating, represented by the blue avatar, with their rating for their *friend* and the *canonical* individual). Subjective confidence was represented by the standard deviation for each rating (e.g., curve width in the illustrations), where lower confidence entails higher standard deviations, and helped account for the influence of memory on choice behavior. *D.* Anchor rescaling. Illustration of how the stranger’s rating is inferred by mentally rescaling the anchor individual’s rating from its perceived relative position on the screen (5), to its absolute position (participant’s rating) on the preference scale.

In summary, training subjects to think of absolutely coded personal preferences in the form of a mental number line allowed us to probe different everyday scenarios at various levels of discriminability and use a well-characterized signal detection model of decision-making (Tanner & Swets, 1954). Discriminability was determined by the absolute distance between individuals on a scale and how confident subjects were about a particular preference. Furthermore, this modeling approach allowed us to account for differing mnemonic demands related to the subject knowing their ratings better for themselves (*self*) than their *friend* and *canonical* ratings (Fig. 1C). Casting our paradigm in terms of coordinate transforms, we investigated how personal preferences are represented in the brain in two complementary ways. First, how the stranger’s rating is inferred by mentally rescaling the *anchor* individual’s rating from its perceived relative position on the screen, to its absolute position (the participant’s actual rating) on the preference scale (i.e., anchor rescaling). Second, uncertainty in preference discrimination based on the absolute distance between the stranger’s rating and the ratings of the personally familiar individuals that the stranger is compared with (i.e., choice discriminability). We subsequently refer to these two elements of representing personal preferences in our task as anchor rescaling(Fig. 1D) and choice discriminability(Fig. 1C), respectively.

## Results

### Behavior

Subjects’ ratings of personally known individuals followed a consistent structure. As expected, subjects tended to rate the *canonical* individual towards the middle of scale, 5, with the *self* and *friend* ratings being more evenly dispersed between 1-9 (Fig. 2A-source Data 1). Subjects’ ratings for the *canonical* exemplar and their *friend* were generally consistent before and after scanning; after scanning, subjects were on average within ±1 of their original ratings for 67.7% (SD=8.15%) of trials and only made large deviations from their original ratings >=±3 on 8.54% (SD=6.2%) of occasions (Fig. 2B-source data 2). Notably, there was a significant effect of *anchor* condition (*self, canonical*, or *friend*) for the average absolute distance between the choice options and the stranger (F(2,22)=10.1;p=.001), where *self* (mean=2.34; SD=.412) and *friend* (mean=2.26; SD=.371) *anchor* conditions had significantly smaller absolute distances than the *canonical* (mean=2.62; SD=.473) *anchor* condition (Fig. 2C-source data 3). Additionally, there were overlapping ratings (the same rating for two individuals on the same scenario) for a small subset of preferences, and there was a significant effect of *anchor* condition in the number of overlapping ratings (F(2,22)=6.74; p=.005), such that there was higher overlap between *self* and *friend* ratings (the *canonical anchor* condition) than between other pairings (Fig. 2-Figure Supplement 1-source data 7). Crucially, trials with overlapping ratings were always considered correctly answered and not included in subsequent behavioral and fMRI analyses. When relating subjective confidence to rating consistency, confidence ratings significantly correlated with rating consistency (t(23)=−4.11;p<.001), where higher confidence ratings had more rating consistency. These results suggest a metacognitive validity of the subjective ratings, relating to memory for specific personal preferences.

During the fMRI experiment, subjects made 72.6% of decisions correctly (SD=1.13%; n=24), after eliminating trials for which answer was correct (i.e., same or equidistant ratings). The mean reaction time (RT) was 4.07 s (SD=.803 s). There was a significant effect of condition (p<.05) for RT (F(2,22)=4.57;*P*=.022; Fig. 2D-source data 4), but not performance (F(2,22)=2.73; *P*=.087). The RT effect was driven by significantly higher RT for *self* versus *friend anchor* conditions (t(23)=2.71;*P*=.012; Fig. 2D). Notably, there was a negative correlation between trial by trial RT and accuracy (t(23)=−5.51;p<.001), i.e., quicker RT for accurate choices. In parallel, RT and rating consistency were also correlated (t(23)=2.55; p=.015), where slower RT related to trials containing inconsistently rated preferences (i.e., poorly remembered preferences),

We then related RT and accuracy to the absolute distance between the stranger’s rating and the ratings of the non-anchor individuals presented as choice options for each trial (i.e., choice discriminability without the use of a model and confidence ratings). As expected, we found that this distance negatively correlated with RT (t(23)=−5.42;p<.001) and positively correlated with accuracy (t(23)=15.8;p<.001); i.e., larger distances related to quicker and more accurate decisions. Highlighting how the metric nature of these absolute distances influenced behavior, we observed a steady increase in the quickness (Fig. 2-Figure Supplement 2-source data 8) and accuracy (Fig. 2E-source data 5) of the participant’s responses when the stranger’s rating was closer to one choice option versus another. Further relating these absolute distances to subjects’ behavior, we then used a signal detection model fit to subjects’ performance that was based on absolute rating distances and subjective confidence ratings. Given the relationship between subjective confidence and rating consistency, subjective confidence ratings also helped account for the different mnemonic demands induced by the different conditions.

#### A computational model of latent preferences

We quantified the difficulty of discriminating a particular preference by fitting a formal signal detection model (Tanner & Swets, 1954) based on the absolute distance between the two choice (*non-anchor*) individuals on a scale and subjective confidence. Specifically, we characterized discriminability using the entropy of decision probabilities based on a softmax function of likelihood and confidence ratings (Fig. 1C). Using this modeling approach allowed us to characterize the absolute distance between the ratings, along with the subjects’ confidence ratings, in a single measurement of trial-specific choice discriminability. Importantly, high entropy corresponds to lower choice discriminability that is induced by similar ratings. To estimate the requisite softmax (sensitivity or precision) parameter, we modeled performance in terms of entropy (*H*) over trials (and subjects) using a simple linear regression model. The ensuing behavioral model provided an estimate of the precision parameter (*β*) and associated measure of trial-specific choice discriminability for each subject.

In detail, we want to score the difficulty of each trial. This difficulty is the entropy (*H*) of the choice probability (*p*), based upon a softmax function of the log odds ratio of a target (*stranger*) preference (*T*) being sampled from the distributions of (*non-anchor*) reference subjects A and B. Let the preference of individual *i* have a mean *μ*_*i*_ and standard deviation *σ*_*i*_ then the log probability of sampling *T*, from the preference distribution of the i-th reference subject is (ignoring constants):

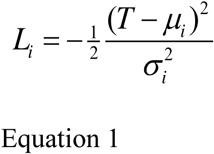

If we assume a precision of *β*, then the probability of choosing subject A over B is a softmax function of the log odds ratio:

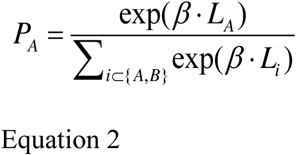

and the difficulty is the entropy:

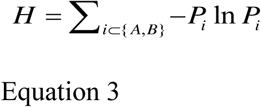

This will give difficulties between 0 and ln(2) i.e., 0.6931. Given the mean and standard deviation reported by subjects, we can evaluate the probability of choosing each (*non-anchor*) individual on each trial, given the *stranger’s* target preference. We used the Bayesian information criterion (BIC) (Schwarz, 1978) to determine if a signal detection model including subjective confidence ratings – as the standard deviation – outperforms a distance-only model. Notably, we found that the model including confidence ratings (BIC=−39,056) outperformed a simpler distance-only model (BIC=− 38,387), where the lower BIC signifies the greater model evidence. In short, this provides strong evidence for a model based on confidence ratings.

This model of choice behavior provided trial-specific measures of choice uncertainty (*H*) that enabled us to identify its fMRI correlates. Following previous work (Daw et al., 2006; Glascher et al., 2010; Daw, 2011), we used the mean precision parameter (*β)*, evaluated over subjects to compute trial-specific choice entropies as a predictor for our fMRI responses. We found that choice entropy negatively correlated with performance over trials (t=−10.9; p <.001). As might be expected, we observed a significant effect of condition for mean choice entropy (F(2,22)=17.6;P<.001; see Fig.2F-source data 6), where choice entropy was significantly higher for *self* than *canonical* (t(23)=6.06;*P*<.001) or *friend* (t(23)=4.23;p<.001) anchor trials (Fig. 2D). This was likely due to lower confidence (i.e., memory demands) for *friend* and *canonical* ratings, relative to ratings of *self* preferences, as well as the smaller mean absolute distances in *self anchor* trials in relation to *canonical anchor trials*.

**Figure 2.**
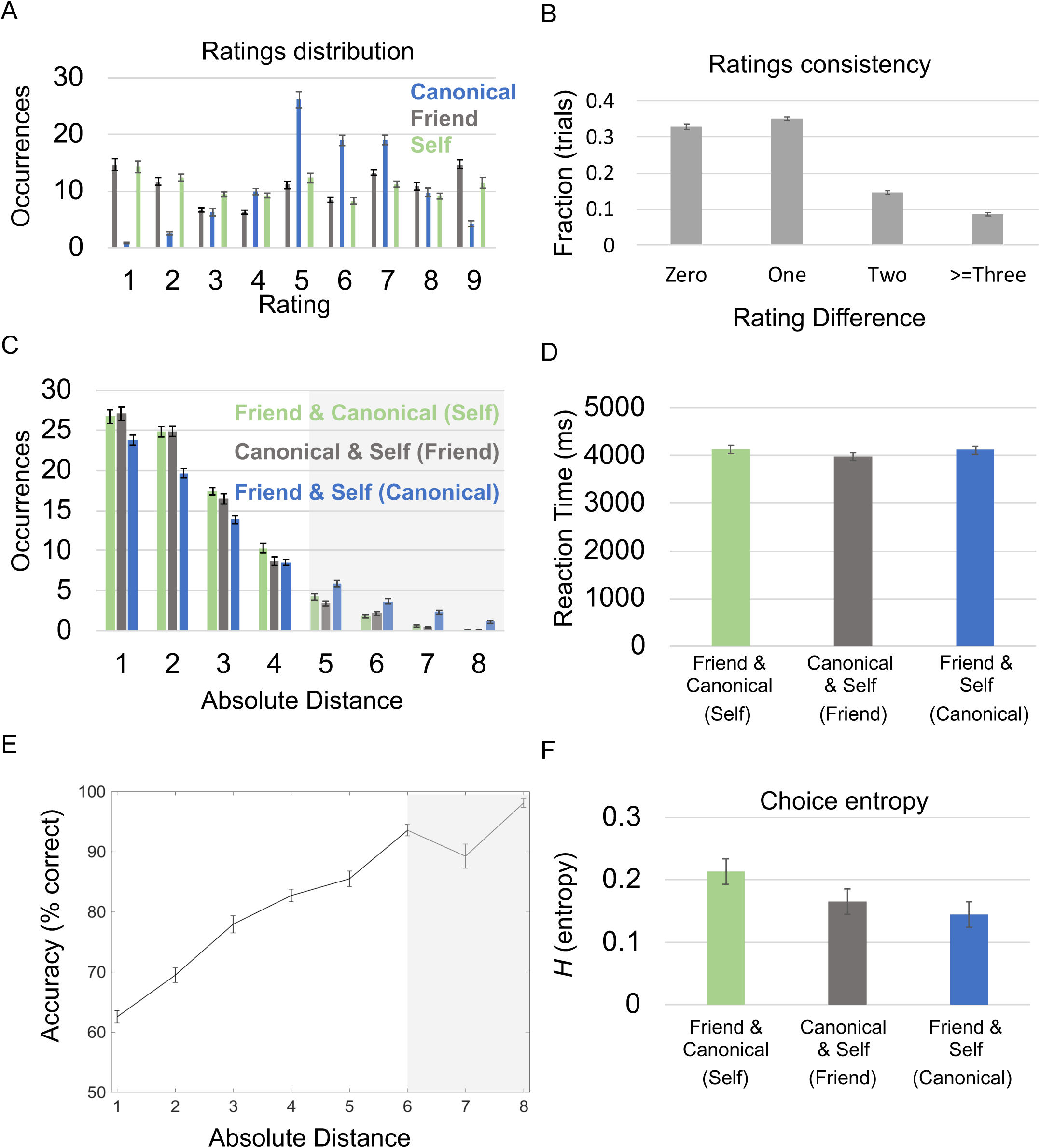
Behavioral Results. *A.* Mean ratings across subjects for each familiar individual and every scenario. Occurrences are out of the 100 total trials per condition. *B.* Rating Consistency. Difference in ratings pre- and post-fMRI scanning for *friend* and *canonical* individuals *C.* Distribution of absolute distances by condition. Mean amount of trials for each *anchor* and absolute distance. Significant effect of condition for absolute distance between the two individuals’ ratings and the stranger’s (p=0.001). Part of the x-axis is shaded in gray because only a fraction of subjects had trials with absolute distances of 5 (*self*: 20/24; *friend*: 19/24; *canonical*: 24/24), 6 (*self*: 14/24; *friend*: 16/24; *canonical*: 23/24), 7 (*self*: 6/24; *friend*: 7/24; *canonical*: 16/24), and 8 (*self*: 3/24; *friend*: 1/24; *canonical*: 13/24, respectively for each *anchor* condition. Individuals being compared are listed in the key with the corresponding anchor/condition name listed in parentheses. *D.* Reaction time: Significant effect of condition for reaction time/decision speed (p=.022). Individuals being compared are listed below each bar with the corresponding anchor condition listed in parentheses. *E.* Absolute distances and performance: Significant relationship (p<.001) between accuracy and the absolute distance between strangers’ ratings and the non-anchor individuals for each trial. 50% represents chance level of accuracy. 7 & 8 on the x-axis are shaded in gray because 18/24 and 13/24 of subjects had trials with absolute distances of 7 & 8, respectively. *F.* Choice entropy: Significant effect of condition for choice entropy (p<0.001). Individuals being compared listed below each bar, with corresponding anchor listed in parentheses. All error bars showing mean ± SEM. For absolute distance plots, the absolute value of each absolute distance was rounded to the closest integer and plotted from 1 to 8.

### fMRI Analyses

#### Choice Discrimination

To test how strangers’ preferences are compared to personally known people, we characterized the brain’s response to uncertainty in preference discrimination. Specifically, we were interested in what brain regions related to choice discrimination in different reference frames.

In a whole-brain analysis – comparing differences in entropy effects between the three conditions (*self, friend, canonical*) – the only significant effect of condition, and the strongest effect of any area in the brain for this contrast, was in a right entorhinal/subicular area (x=24,y=−28,z=−22; F-stat:9.87; Z-score=3.46; SVC peak-voxel p=.034; Fig. 3A-B), extending into anterior parahippocampal cortex. In other words, the right entorhinal/subicular effect of discriminability depended upon the *anchor* condition.

**Figure 3.**
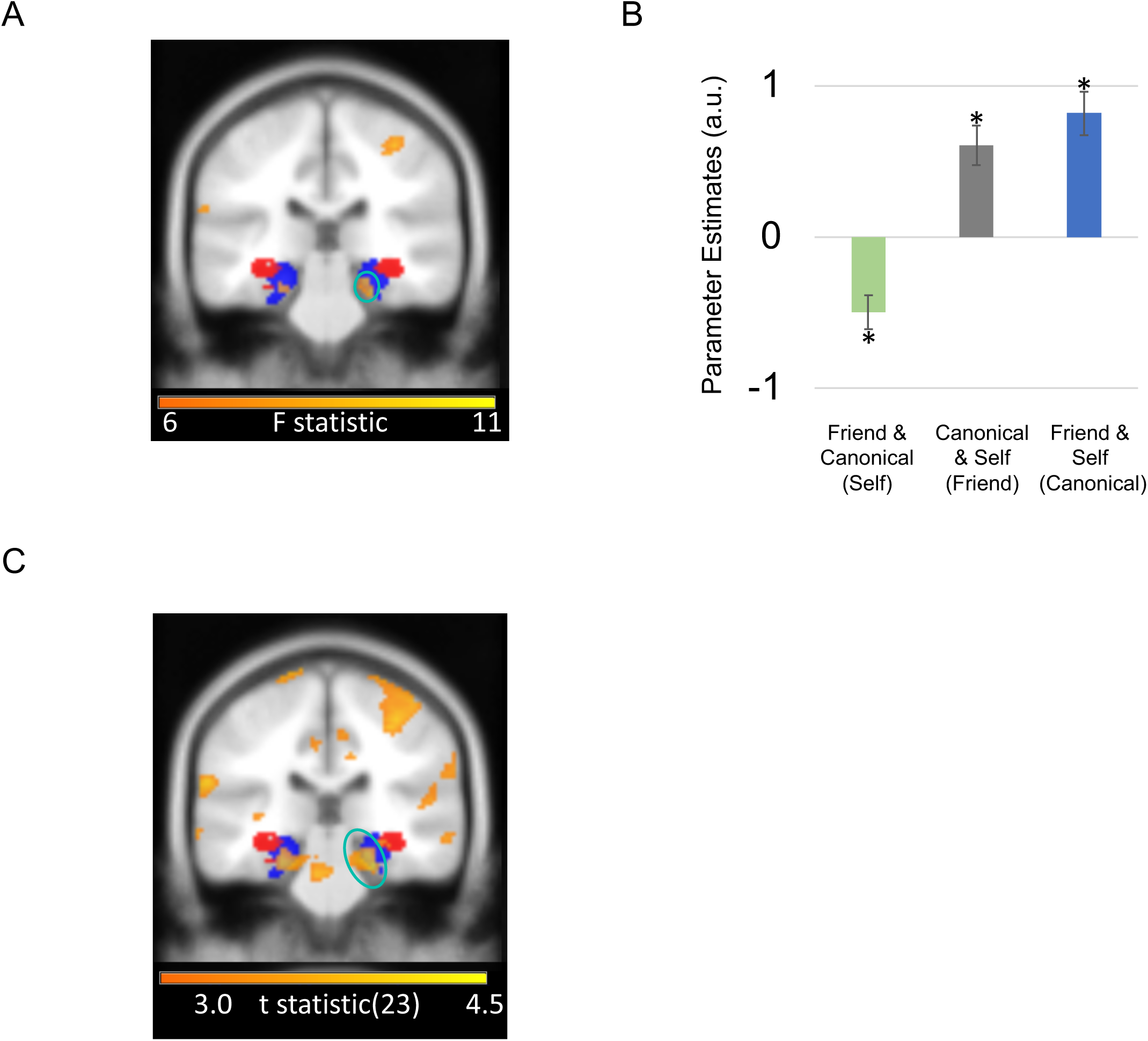
Entorhinal/Subicular Choice Discrimination Effects. *A.* Coronal image of right entorhinal/subicular region exhibiting an effect of choice discriminability/entropy by condition circled in turquoise. B.: Effect size for a 10-mm sphere around right entorhinal/subicular region exhibiting effect of choice entropy by condition (mean ± SEM). Asterisk marks significance at p<.05. C. Coronal image showing right entorhinal/subicular (circled in turquoise) correlation with choice entropy for f*riend* and *canonical anchor* trials (i.e., choices involving self-comparisons) versus *self anchor* trials (i.e., trials comparing canonical versus friend ratings). Portion of left entorhinal/subicular region showing same effect is also visible. A positive effect size indicates a positive BOLD correlation with choice entropy (i.e., ambiguous choices), whereas a negative effect size indicates a negative BOLD correlation with choice entropy in the same comparison (i.e., straightforward choices). All highlighted regions survived FWE correction for multiple comparisons at p< 0.05 and are displayed at an uncorrected statistical threshold of p<.005 for display purposes. For visualization purposes, entorhinal/subicular (blue) and hippocampal body (red) probabilistic masks from the Jülich SPM Anatomy toolbox are presented (Amunts et al., 2005).

Subsequent t-tests on the entorhinal/subicular region exhibiting choice discriminability/entropy effects indicated that there was a significant positive correlation with entropy (i.e., less discriminable/ambiguous choices) for *canonical* (t(23)=2.75;p=.011) and *friend* (t(23)=2.36;p=.027) *anchor* trials, while there was a significant negative correlation with entropy (i.e., more discriminable/straightforward choices) for *self anchor* trials (t(23)=−2.39;p=.025; Fig. 3B). Contrasting *canonical* and *friend* versus *self anchor* trials, we observed that the right entorhinal/subicular effect (right: Z-score=4.02; peak voxel p=.004 Fig. 3C) was the strongest effect of any area in the brain for this contrast. Note that *canonical* and *friend* anchor trials involved choices with *self* preferences. Subjects either decided whether a *stranger’s* rating for a scenario was closer to the *self* versus *canonical* individual (*friend anchor*), or the *self* versus the *friend* (*canonical anchor*). Conversely, *self anchor* trials involved deciding whether a *stranger’s* rating was closer to the *friend* versus the *canonical* individual. In other words, the entorhinal/subicular region responded to more fine-grained/ambiguous choices involving self-comparisons, but readily discriminable choices otherwise.

We then investigated which brain regions responded to choice entropy (i.e., how discriminable the choices were). However, we did not find any significant regions responding to increasing (i.e., a positive correlation between the fMRI signal and choice entropy/less discriminable choices) or decreasing choice entropy (i.e., a negative correlation between the fMRI signal and choice entropy/more discriminable choices) anywhere in the brain.

#### Anchor Rescaling

In our task, reference frame transformations rely on flexibly relating different known individuals’ ratings to each other. We wanted to capture the initial demands of mentally rescaling the anchor and, consequently, the corresponding stranger from its observed (relative) position on the screen to its actual rating (absolute position) on the scale. To test this, we investigated which regions responded to how far the *anchor’s* preference deviated from the middle of the scale (Anchor Rescaling; |anchor rating-5|). In a whole-brain analysis, the only significant main effect of increased anchor rescaling towards the limits of the scale was in the superior parietal lobule (SPL) bilaterally (left: x=−57,y=−52,z=44; Z-score=4.09; FWE cluster p=.024; right: x=33,y=−64,z=54; Z-score=3.99; FWE cluster p=.005; Fig. 4-Figure Supplement 1).

However, we did not observe any significant effects for decreased anchor rescaling (maintaining the rating in the middle of the scale). Likewise, we did not observe any regions showing an interaction with anchor condition.

#### Accuracy

Further characterizing the functional contribution of different brain regions, we asked if regional responses during memory-guided decisions depended on whether subjects made a correct, or incorrect choice for each trial. In a whole-brain analysis, we observed significant activation for correct trials in the right ventral striatum (x=24,y=8,z=−6; Z-score=4.69; FWE cluster p<.001; Fig. 4A-B) and left posterior parietal cortex (PPC) in the depth of intraparietal sulcus (x=−24,y=−64,z=42; Z-score=4.30; FWE cluster p=.005; Fig. 4-Figure Supplement 2). We did not observe any significant activation that preceded incorrect choices, nor significant interactions between anchor condition and accuracy.

**Figure 4.**
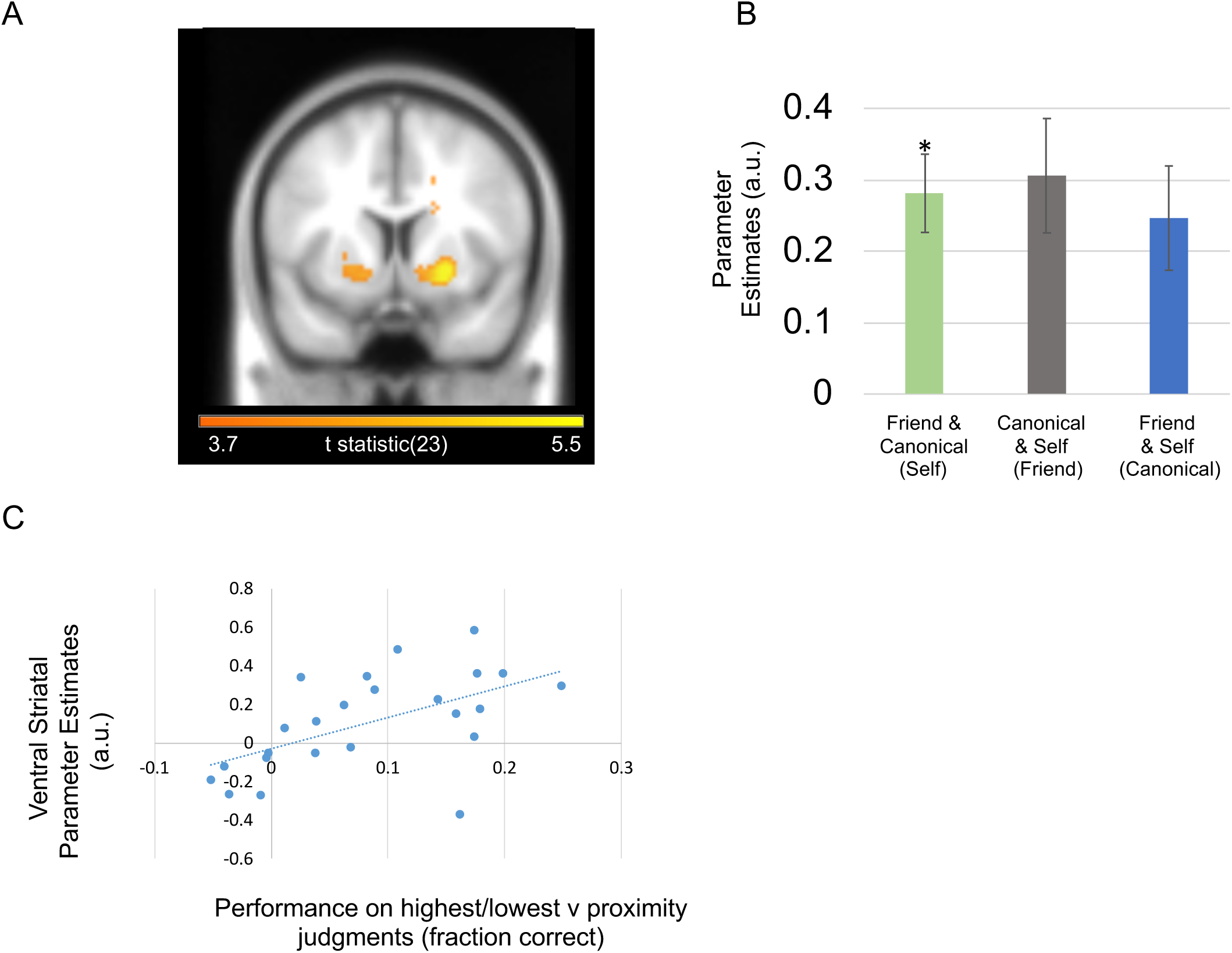
Striatum and Decision Accuracy. *A.* Ventral striatal activity related to correct versus incorrect choices. The ventral striatal cluster survived cluster-level FWE correction at p< 0.05 and is displayed at an uncorrected statistical threshold of p<.001. *B.* Effect sizes for a 10-mm sphere around right ventral striatum peak voxel (mean ± SEM). Individuals being compared list below with anchor in parentheses. A positive effect size indicates a positive BOLD correlation with correct choices. Asterisk marks significance at p<.05. *C.* Plot showing between-subject correlation for subjects’ ventral striatal fMRI signals with behavioral performance. Ventral striatal effects were for highest/lowest judgments versus proximity judgment trials (GLM2). Behavioral performance was taken from subjects’ performance on trials, where highest/lowest judgments could be used, versus their performance on trials where a proximity judgment could be used. Subjects who performed better for highest/lowest judgments exhibited increased ventral striatal activity for the same contrast.

Finally, to test whether the right ventral striatum accuracy effect was driven by choices where highest/lowest judgments between the two choice individuals needed to be made, we split decision-making trials in terms of whether a choice required determining which choice option was highest/lowest, instead of closest. Ignoring the three anchor conditions, we split the data into two conditions; one where the target was between the two choice individuals, and the second where the target was between either higher or lower than both choice individuals. We asked whether subjects who performed better for the latter scenario, also recruited the ventral striatum when making highest/lowest versus proximity judgments. We found a between-subject correlation between our right ventral striatal effect for highest/lowest judgments and subject performance for highest/lowest versus proximity judgments (r=.561;p=.004) (Fig. 4C).

#### Remaining fMRI analyses

Asking whether the time spent deliberating and trying to remember a particular choice option’s preferences influenced any neural responses during our task, we investigated whether any fMRI signals during the social decision-making task correlated with RT. We observed signals in the posterior parietal cortex/precuneus (left: x=−18,y=−58,z=44; Z-score=4.49; FWE cluster p<.001; right: x=30,y=−73,z=42; Z-score=4.07; FWE cluster p=.001; Fig. 4-Figure supplement 3), left motor cortex (x=−33,y=5,z=48; Z-score=4.35; FWE cluster p=.033), and lateral occipital cortex (left: x=−24,y=−97,z=−6; Z-score=4.26; FWE cluster p<.001; right: x=27,y=−91,z=4; Z-score=4.35; FWE cluster p=.033) that positively correlated with RT. However, we observed no significant negative correlation with RT anywhere in the brain, nor significant interactions between *anchor condition* and RT. Lastly, we also tested whether there were neural responses specifically linked to participants’ subjective confidence ratings for a given choice and did not observe significant responses in any brain region.

## Discussion

Using fMRI, a signal detection model, and a novel memory-guided decision-making paradigm, we asked how different brain regions integrate relatively coded knowledge of different people’s personal attributes within absolutely coded metric spaces (Fig. 1). Focusing on reference frame-sensitive responses, we identified an entorhinal/subicular region that related to the absolute distance between the stranger’s rating and the ratings of choice options (Fig. 3). In parallel, superior parietal lobule activity increased when the *anchor* required a mental shift from the middle of the scale towards the periphery(Fig. 4-Figure supplement 1). Finally, we found that striatal responses also preceded correct choices, which were partially driven by decisions about which individual had the highest or lowest preference rating (Fig. 4). In what follows, we relate these memory-guided choice findings to the wider literature – and to the different hippocampal formation responses we observed. We then speculate on potential neural computations induced by our task.

### Entorhinal-subicular representations of social knowledge along multiple scales

We observed entorhinal/subicular responses to more precise, fine-grained choices, if self-comparisons were involved, but more straightforward choices otherwise. This result is partially a consequence of lower confidence when rating the *friend* and *canonical* individual, as well as smaller mean absolute distances between those two individuals and the stranger, which induced significantly higher choice entropy for the *self anchor* condition; i.e., a more ambiguous/less discriminable choice scenario. Despite the difference between the *anchor* conditions in mean choice entropy (potentially due to an entropy ceiling effect in the *self anchor* condition), a similar distinction was not observed in the hippocampal body, or other brain regions. Notably, in rodents, hippocampal place representations are typically self-referenced and continuous (Buzsaki & Moser, 2013). In contrast, entorhinal/subicular grid, as well as object and boundary vector representations use multiple reference frames, and can either be continuously or discretely coded (Hartley et al., 2013; Diehl et al., 2017; Høydal et al., 2018). This highlights a potential functional dissociation to memory-guided decision-making involving different reference frames, where the body of the hippocampus could relate to self-referenced sampling of prior experience (Ezzyat & Davachi, 2014; Bornstein & Norman, 2017; Bakkour et al., 2018; Shadlen & Shohamy, 2016). While in contrast, we observe entorhinal/subicular sensitivity to the depth of personal knowledge for different familiar individuals (i.e., how well we know someone’s personal preferences). Further support for this distinction stems from the putative role of the entorhinal cortex (Buzsaki & Moser, 2013; Maas et al., 2015; Navarro et al., 2015) and subicular region (Dalton & Maguire, 2017) in integrating incoming sensory input with learned hippocampal representations. Building on recent studies of macaque and human entorhinal cortex in spatial and non-spatial tasks, our results potentially speak to the organizational principles governing metrically coded maps (Doeller et al., 2010; Chadwick et al., 2015; Constantinescu et al., 2016; Lositsky et al., 2016; Vass et al., 2017; Julian et al., 2018; Meister et al., 2018; Nau et al., 2018), along with the implicit encoding of discrete graphs (Garvert et al., 2017), which may be located within the same brain region. This architecture portends a domain general role for entorhinal cortex in memory-guided decision-making involving multiple reference frames.

### Parietal rescaling of metrically coded preferences

In our task, participants retrieve and non-linearly rescale different mental number lines of personal preferences. Following previous observations of posterior parietal involvement in maintaining a mental number line (Dehaene et al., 1993, 2003) and manipulating the position of items in spatial working memory (Koenigs et al., 2009), we observe bilateral superior parietal lobule responses during trials when participants need to mentally rescale the anchor’s relative position in the center towards an absolute position on the periphery of the scale. Moreover, our observation of superior parietal lobule involvement in mental rescaling of metrically scaled preferences meshes well with the putative notion that parietal areas track magnitudes (Bueti & Walsh, 2009).

### Striatal involvement in knowledge-guided social decision-making

We observed a ventral striatal signal corresponding to correct choices, which also related to choices where highest/lowest–as opposed to proximity–judgments could be made. This distinction between hippocampal-entorhinal versus striatal responses to different knowledge-guided decision-making strategies closely follows spatial navigation strategies; where the subjects align themselves to a single landmark, instead of a boundary, in order to infer the direction of a goal location (Doeller & Burgess, 2008). The implicit functional neuroanatomy may parallel the dissociation between hippocampal and striatal responses observed in the current study. This follows since boundary-oriented navigation is linked to the hippocampus, whereas landmark-based navigation is related to the striatum (Doeller et al., 2008). More generally, this striatal versus hippocampal dissociation highlights a potential mechanism for how reinforcement/procedural learning could reduce metrically-coded relational knowledge into more efficiently usable heuristics (Simon & Daw, 2011; Gershman & Daw, 2017), paralleling categorical versus coordinate-based judgments in spatial cognition (Kosslyn, 1987; Baumann & Mattingley, 2014).

### Allocentricity and the dimensionality of preferences

Our findings highlight the role of the hippocampal formation, namely the entorhinal cortex and subiculum, in the functional anatomy of how we transform relative and absolute reference frames during decision-making. However, since all of our conditions involve a transformation, it is unclear whether humans ever preferentially use an absolute or ‘allocentric’ frame of reference during decision-making without drawing upon their own frame of reference. Even in spatial and episodic memory this question is difficult to resolve (see Filimon, 2015); since self-referencing helps us relate past experience to our current environment (Vann et al., 2009). Useful clues about how we flexibly transform relative and absolute reference frames come from the social psychology literature. A subset of social psychology has focused on how individualized representations of others’ preferences are generated by anchoring to a known preference, and then adjusting accordingly, a phenomenon known as ‘anchoring and adjustment’ (Epley & Gilovich, 2001; Epley et al., 2004; Tamir & Mitchell, 2013). Given their similarities, jointly studying ‘anchoring and adjustment’ with boundary-oriented navigation may offer a promising avenue for translational research across species and levels of analysis.

We tested metrically-coded preferences along a single dimension (namely the likelihood of people preferring things), but the true dimensionality of personal preferences is less clear. It is plausible that we learn about others’ preferences within a multi-dimensional trait space (Tamir & Thornton, 2018). Yet we do not know the number of dimensions that support this trait space (Tavares et al., 2015; Bellmund et al., 2018). Further work may help relate what we know about navigating physical space with the mental exploration of more abstract spaces (Aronov et al., 2017; Kaplan et al., 2017a).

Our study induces a spatial (metric) strategy, preventing us from asking whether humans must use map-like coding in order to relate others’ preferences to their own. Furthermore, decisions in our task involve an abstract one-dimensional spatial discrimination, where subjects determine the relative proximity of a stranger’s rating to the non-anchor individuals. Future work can implicitly test discrimination of metrically coded decision variables in different reference frames more implicitly (i.e., without an explicit spatial element) to determine whether the spatial element is required for hippocampal-entorhinal involvement. Still, theoretical work has provided support for the use of a spatial strategy, where non-human primate research has investigated social cognition in terms of coordinate transforms (Chang et al., 2013). Social coordinate transformation experiments have captured how social variables are encoded in individual neurons – and how encoding changes over different computational stages during social decision-making (Chang, 2013), like current versus long-term frames of reference (Boorman et al., 2013). Extending such work to abstract knowledge can potentially help us understand how coordinate transformations can generally guide everyday decisions.

### Conclusion

Metric coding of decision variables informs decisions by providing coordinates and boundaries that can be translated between different frames of reference. We provide evidence that neural computations – that integrate relative and absolute coordinates during spatial navigation – also extend to relating others’ personal attributes to our own. Consequently, these data provide important clues about how hippocampal-entorhinal map-like coding may facilitate memory-guided decision-making in a domain general manner.

## Materials and Methods

### Subjects

Twenty-four healthy adult subjects were studied and compensated (16 female; mean age in 25.5 y; SD of 5.38 y) and gave informed written consent to participate. This study was approved by the local research ethics committee at University College London. The study was conducted in accordance with Declaration of Helsinki protocols. All subjects were right-handed had normal or corrected-to-normal vision and reported good health with no prior history of neurological disease.

### Task

Stimuli were presented using the Cogent (http://www.vislab.ucl.ac.uk/cogent.php) toolbox running in MATLAB (Mathworks, Natick, MA, USA). Subjects performed a self-paced (max 9s), forced-choice, social decision making-task featuring 100 different personal preferences for 3 personally familiar individuals, which included themselves. Subjects viewed a personal preference for a novel stranger, represented by an avatar (Fig. 1), that was presented on a scale relative to one of three known individuals (*anchor*). Immediately prior to scanning, subjects were first trained to infer the stranger’s rating and then look below the scale in order to decide which of the two remaining familiar individuals was closer to that rating. All of the information needed to perform the task was presented at once on the screen and the only aspects of the task that varied from trial to trial were the anchor and choice options.

All ratings were obtained from subjects approximately 45 minutes prior to fMRI scanning. Subjects gave likelihood ratings about 110 everyday scenarios (e.g. eating spicy food; see Figure 1-Source data 1 for all scenarios) from 1-9 on a 0-10 scale (0 being impossible to 10 being extremely likely) for themselves and two other familiar individuals. For one of the familiar individuals (*friend*), subjects were asked to choose their closest friend of the same gender that had the most distinct personality from the subject and – unlike the subject – ideally lived outside of London, in order to induce different ratings between individuals. The third individual was a typical/*canonical* individual of the same gender named Mr/Ms. Normal. Mr/Ms. Normal was supposed to correspond with what they thought a normal person/exemplar at their stage of life would do for each scenario (Fig. 1A). During the rating period, subjects were instructed to keep track of how their ratings related to each other for a given scenario (e.g., the participant being a bit more likely to have spicy food than their friend). Crucially, subjects also gave confidence ratings for the *canonical* exemplar and their *friend* on every scenario on a scale from 1-5 (No Idea to Very Sure) in order to assess the consistency of their rating (Fig. 1A). As a second confirmation of rating consistency, subjects also rated the familiar individuals for every scenario after the fMRI task.

Subjects then performed a brief practice version of the fMRI social decision-making task outside of the scanner, where subjects viewed a personal preference for a *stranger* that was presented on a 0-10 scale relative to one of the known individuals (*anchor*). Subjects had a maximum of 9 seconds to decide whether the *stranger’s* rating was closer to the two remaining (*non-anchor*) individuals. The *anchor* was indicated by presenting their initials, which would either be the friend’s initials, ME (Self), or MN, which was an abbreviation for Mr/Ms. Normal (*Canonical*). All information related to the decision was presented at once, with the scenario being listed above the stranger and anchor individual on the scale (Fig. 1B). Directly below the scale “closer to self/friend/canonical individual (A) or self/friend/canonical individual (B)” was written. After making a decision, there was then a jittered intertrial interval (ITI) period (mean=2.43s; range=0.25-9s) where a white fixation point overlaid on a black background was presented.

Crucially, the anchor individual was always placed in the middle of the scale, so that subjects needed to use prior social knowledge in order to infer the *stranger’s* absolute preference and form a mental number line of the *non-anchor* preferences. In other words, subjects had to infer the stranger’s true (absolute) position in relation to the anchor’s rating for that scenario and the closest boundary (0 or 10). For example, if the stranger was ¾ to the right of an anchor individual that was rated a 9, the participant would infer that the stranger’s rating was ¾ of the way between 9 and 10 (the right boundary of the scale). Consequently, the subject would indicate the stranger’s rating would approximately be near 10 (actual rating=9.75). We assumed that subjects represented people’s preferences on this number line, similar to a mental number line (Dehaene et al., 1993), thereby enabling them to choose the *individual* (A or B) that was closest to the *stranger*. If both individuals in the choice were the same or equidistant from the stranger, subjects were instructed that either answer was counted as correct. Same/equidistant trials were not used in either behavioral or fMRI analyses.

Subjects then practiced the task using the last 10 scenarios they rated, which were repeated three times each. On the first few practice trials, subjects verbally rehearsed transforming the anchor’s rating from the middle of the screen to its actual (absolute) position and then inferring the stranger’s rating with verbal feedback from the experimenter until they performed the transformation correctly.

Then during fMRI scanning, subjects performed the task for the remaining 100 scenarios in three different self-paced runs (once for each anchor individual) each lasting approximately 10 minutes (a maximum of 15 minutes). From a relative viewpoint, the stranger was either ±1/4 or 3/4 of the way between the anchor individual and the boundary ratings of 0 and 10, which was presented equally by run and condition. Subjects were instructed that it was a different stranger on each trial, so the stranger didn’t conserve any preferences. To further emphasize the lack of conserved preferences, the stranger avatar randomly alternated between five colors (red, green, yellow, magenta, and cyan). Once again after completing the fMRI task, subjects gave likelihood ratings for the friend and canonical individuals outside of the scanner.

### Computational Model

We used a trial-by-trial computational model of subjects’ performance in order to quantify how subjects’ individual ratings and confidence judgments determined subjects’ choice discriminability during social decision-making (see Fig. 1C). Choice discriminability (or inverse precision) was measured using Shannon entropy (Shannon, 1948). Shannon entropy (*H*) was calculated by estimating the distribution of choice probabilities using a softmax function of individual ratings. This included the standard deviation (*σ*_*i*_) derived from subject’s confidence ratings (*C*_*i*_) for each scenario (*σ*_*i*_ =1 for all self ratings), where the constant term accommodated the assumed stability (reduced memory demands) of self ratings, as well as the dynamic range of confidence ratings.

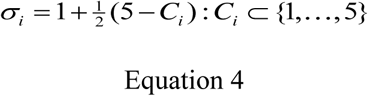

Furthermore, the inverse temperature parameter (*β*) of the softmax function was optimized using each subject’s trial-by-trial performance. We constructed predictions of neuronal responses in terms of the Shannon entropy (*H*) of the choice probability in each trial for fMRI (Kaplan et al., 2017b).

We optimized subject-specific parameters across trials using maximum likelihood estimation and the optimization toolbox in MATLAB (MathWorks, Inc). We then calculated trial-by-trial parameter estimates of *H* (choice discriminability) using the group average softmax (precision or inverse temperature) parameter. The ensuing trial-by-trial choice entropy measures were then used to predict BOLD responses in our neuroimaging analyses.

### fMRI Acquisition

Functional images were acquired on a 3T Siemens Trio scanner. BOLD T2*-weighted functional images were acquired using a gradient-echo EPI pulse sequence acquired obliquely at 45° with the following parameters: repetition time, 3,360 ms; echo time, 30 ms; slice thickness, 2 mm; inter-slice gap, 1 mm; in-plane resolution, 3 × 3 mm; field of view, 64 × 72 mm^2^; 48 slices per volume. A field-map using a double echo FLASH sequence was recorded for distortion correction of the acquired EPI (Weiskopf et al., 2006). After the functional scans, a T1-weighted 3-D MDEFT structural image (1 mm^3^) was acquired to co-register and display the functional data.

### fMRI Analysis

Functional images were processed and analyzed using SPM12 (www.fil.ion.ucl.uk/spm). The first five volumes were discarded to allow for T1 equilibration. Standard preprocessing included bias correction for within-volume signal intensity differences, correction for differences in slice acquisition timing, realignment/unwarping to correct for inter-scan movement, and normalization of the images to an EPI template (specific to our sequence and scanner) that was aligned to the T1 Montreal Neurological Institute (MNI) template. Finally, the normalized functional images were spatially smoothed with an isotropic 8-mm full-width half maximum Gaussian kernel. For the model described below, all regressors, with the exception of six movement parameters of no interest, were convolved with the SPM hemodynamic response function. Data were also high-pass filtered (cut-off period = 128 s). Statistical analyses were performed using a univariate GLM with an event-related experimental design.

#### GLM1

There were two periods of interest, the self-paced (9s maximum) social decision-making and jittered baseline inter-trial interval (ITI), which were modeled as boxcar functions and convolved with a canonical hemodynamic response function (HRF) to create regressors of interest. For each social decision-making regressor (*self, friend*, and *canonical* anchor trials), there were parametric regressors based on accuracy (whether the choice was correctly answered; 1=incorrect choice; 2=correct choice), anchor rescaling, choice entropy (discriminability), reaction time (RT), and subjective confidence ratings. Inferences about these effects were based upon t- and F-tests using the standard summary statistic approach for second level random effects analysis.

#### GLM2

In a follow-up analysis to assay whether the ventral striatum accuracy effect was driven by choices where highest/lowest judgment between the two choice individuals needed to be made, we split decision-making trials in terms of whether a choice required determining which choice option was highest/lowest, instead of closest. Ignoring the three conditions, we split the data into two conditions; one where the target was between the two choice individuals, and the second where the target was either higher or lower than both choice individuals.

All initial analyses were whole-brain analyses. Subsequently, post hoc statistical analyses were conducted using 10-mm radius spheres in MarsBar toolbox (Brett et al., 2002) within SPM12 around the respective peak voxel specified in the GLM analysis. This allowed us to compare the effects of different parametric regressors of interest (e.g., to determine whether an accuracy effect was present in a region defined by an orthogonal main effect of choice discriminability). This ensured we did not make any biased inferences in our post hoc analyses. We used 10-mm radius spheres for post hoc statistical analyses–instead of clusters–since our hippocampal-entorhinal effects were corrected for multiple comparisons at the peak-voxel level.

Given the previously hypothesized role of the entorhinal/subicular region and hippocampus in absolute versus relative coding of environmental cues, we report whether peak-voxels in these regions survive small-volume correction for multiple comparisons (*p* < 0.05) based on bilateral ROIs in the entorhinal/subicular region (mask used in Chadwick et al., 2015; Garvert et al., 2017) and the hippocampus (mask created using Neurosynth, Yarkoni et al., 2011). For all analyses outside of the ROIs, we report activations surviving an uncorrected statistical threshold of *p* = 0.001 and correction for multiple comparisons at the whole-brain level (FWE *p* < 0.05). Coordinates of brain regions are reported in MNI space.

## Acknowledgments

The authors would like to thank Asaf Gilboa, Jochen Michely, Philipp Schwartenbeck, and Geert-Jan Will for helpful discussion, along with Martin Chadwick and Mona Garvert for providing an entorhinal/subicular mask. We thank Jacob Bellmund and Joshua Julian for helpful comments on a previous version of this manuscript. The authors would also like to thank Megan Creasey and Clive Negus for help with scanning and the Wellcome Centre for Human Neuroimaging for providing facilities. This work was funded by a Wellcome Trust grant to KJF (Ref: 088130/Z/09/Z), and a Sir Henry Wellcome Postdoctoral Fellowship awarded to RK (Ref: 101261/Z/13/Z). The funders had no role in study design, data collection and analysis, decision to publish, or preparation of the manuscript. The authors declare no competing interests.

## Figure Supplements

**Figure 2-Figure supplement 1.**
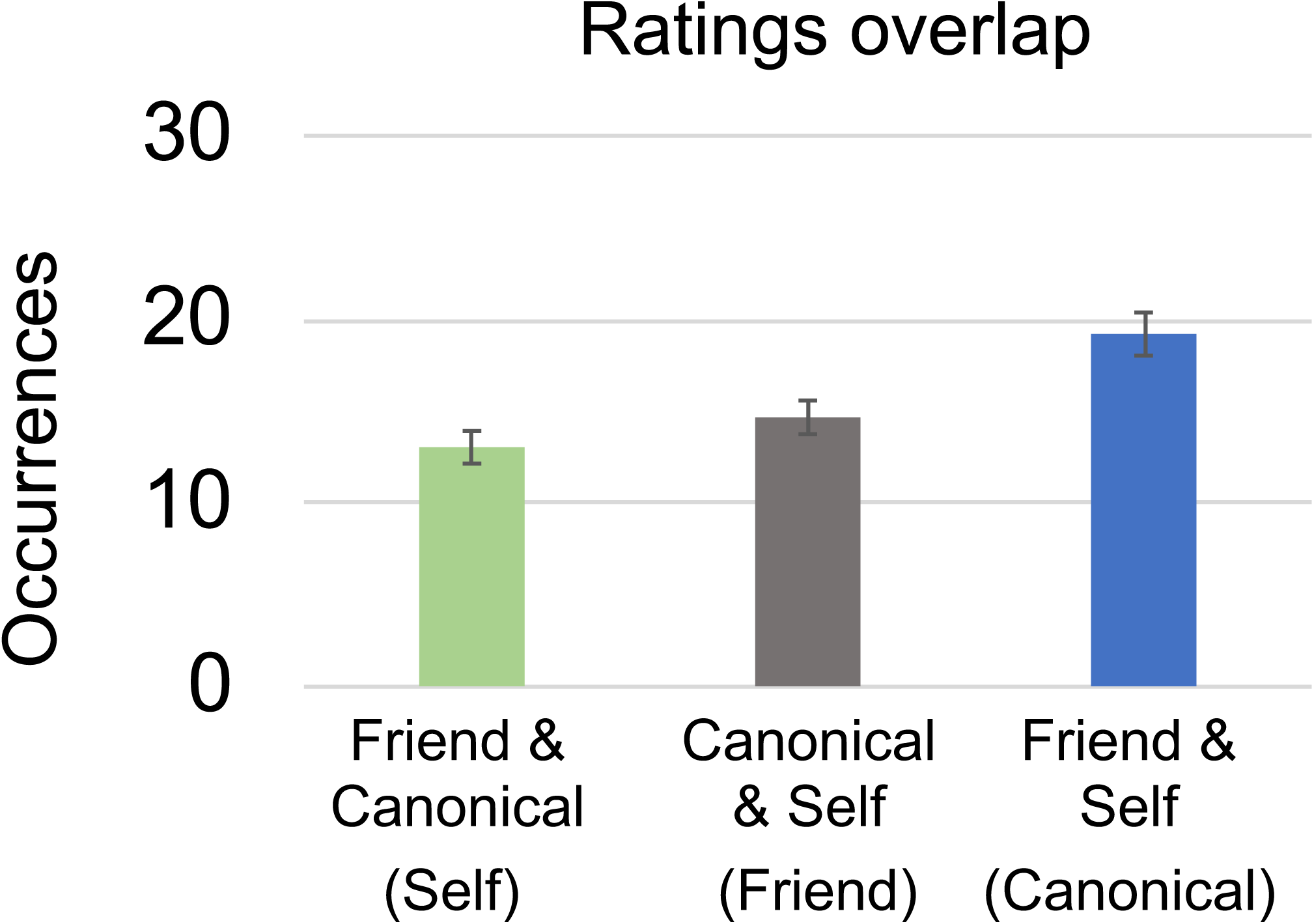
Ratings and Overlap. Significant effect of condition for ratings overlap (the same ratings) between the two individuals (p<0.001). Individuals being compared are listed below each bar with the corresponding anchor/condition name listed in parentheses. Occurrences are out of the 100 trials per condition.

**Figure 2-Figure supplement 2.**
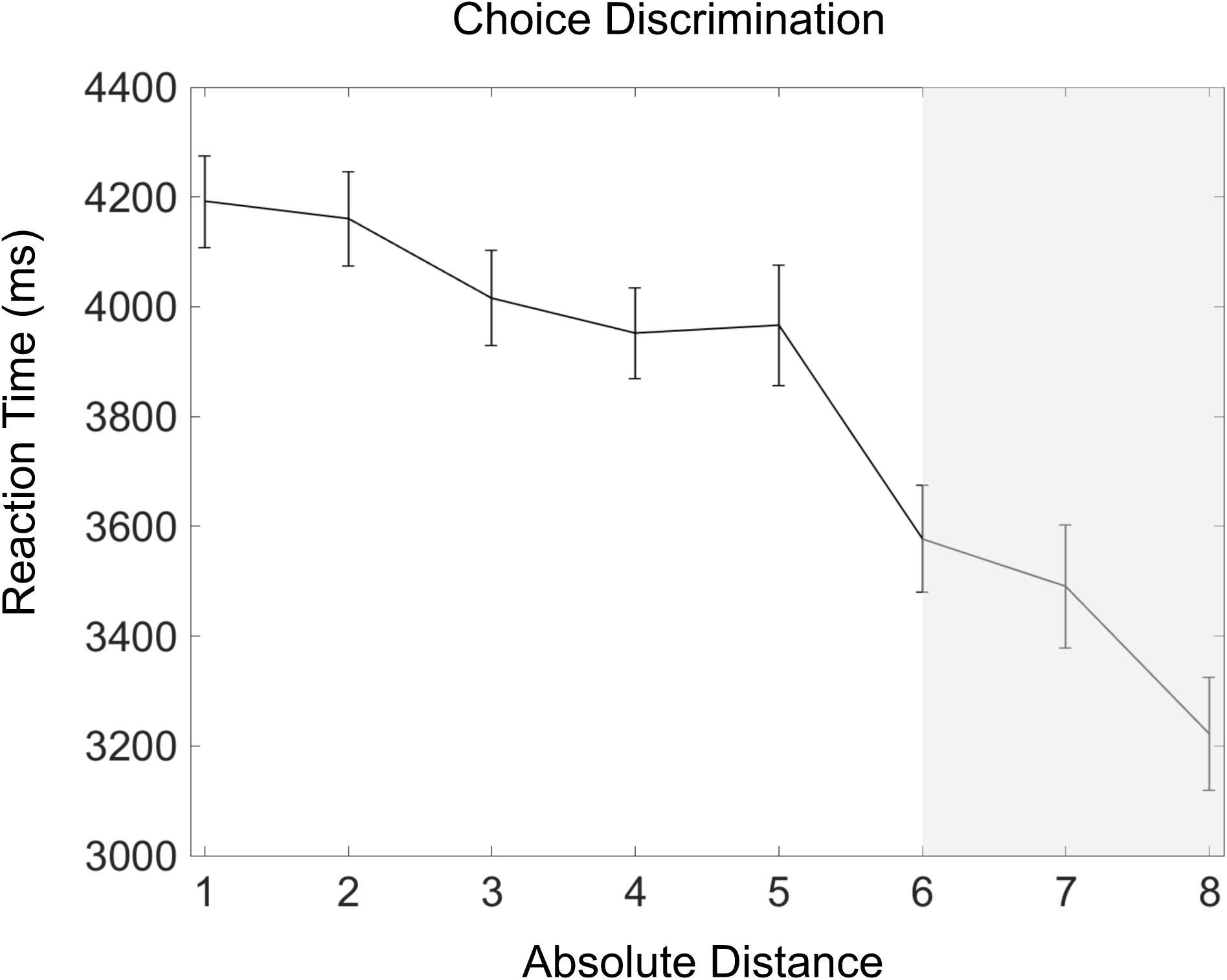
Absolute distances and reaction time: Significant relationship (p<.001) between decision speed (reaction time) and the absolute distance between strangers’ ratings and the non-anchor individuals for each trial. The absolute value of absolute distances were rounded to the closest integer and plotted from 1 to 8. 7 & 8 on the x-axis are shaded in gray, because only 18/24 and 13/24 subjects had trials with absolute distances of 7 & 8 respectively. Error bars showing mean ± SEM.

**Figure 4-Figure supplement 1.**
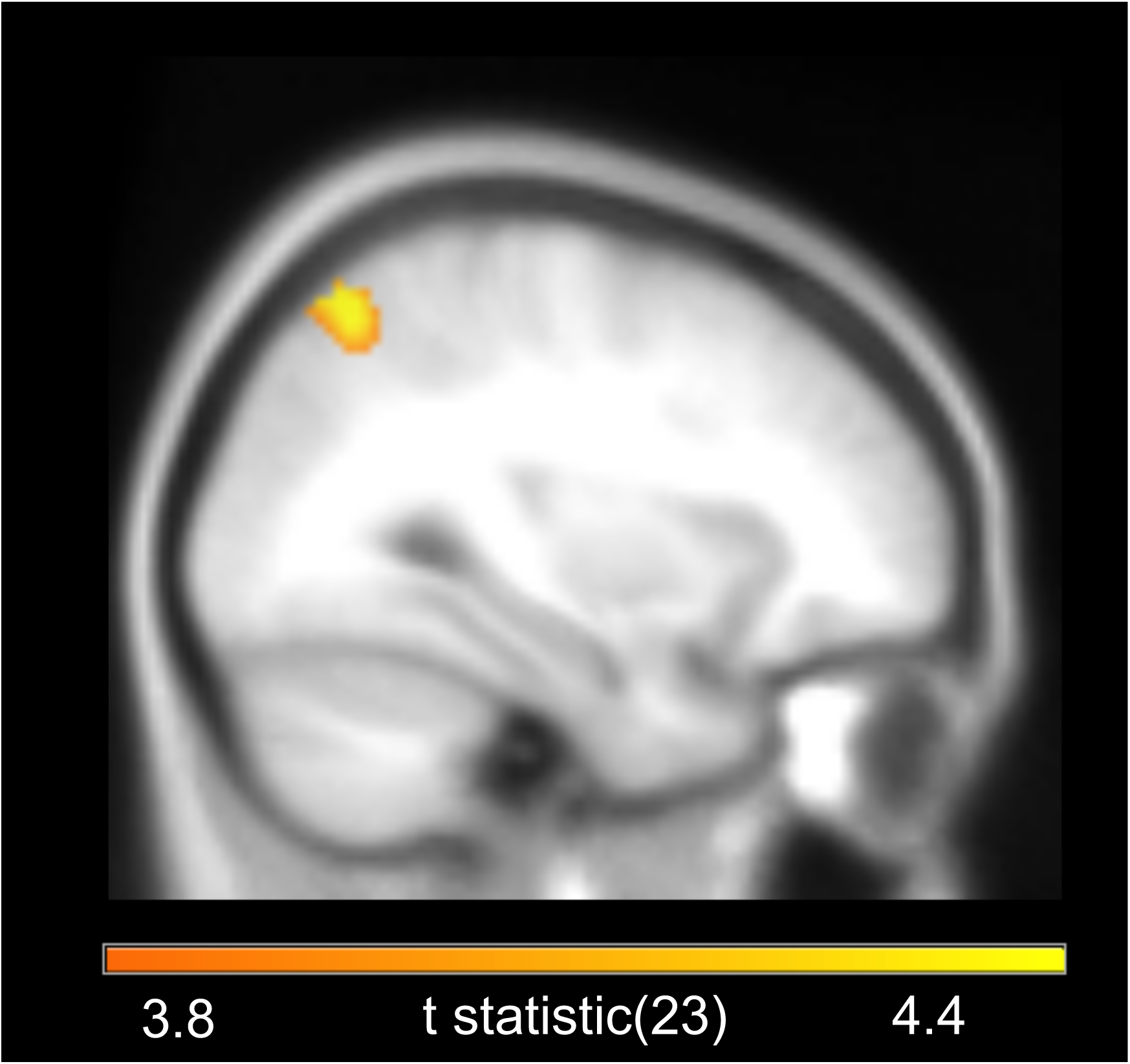
Superior Parietal Lobe Rescaling Effect. Sagittal image showing superior parietal lobule effect of mentally rescaling anchor towards the periphery in either direction, as seen in Figure 1D. Highlighted region survived cluster-level FWE correction at p< 0.05 and image is displayed at an uncorrected statistical threshold of p<.001.

**Figure 4-Figure supplement 2.**
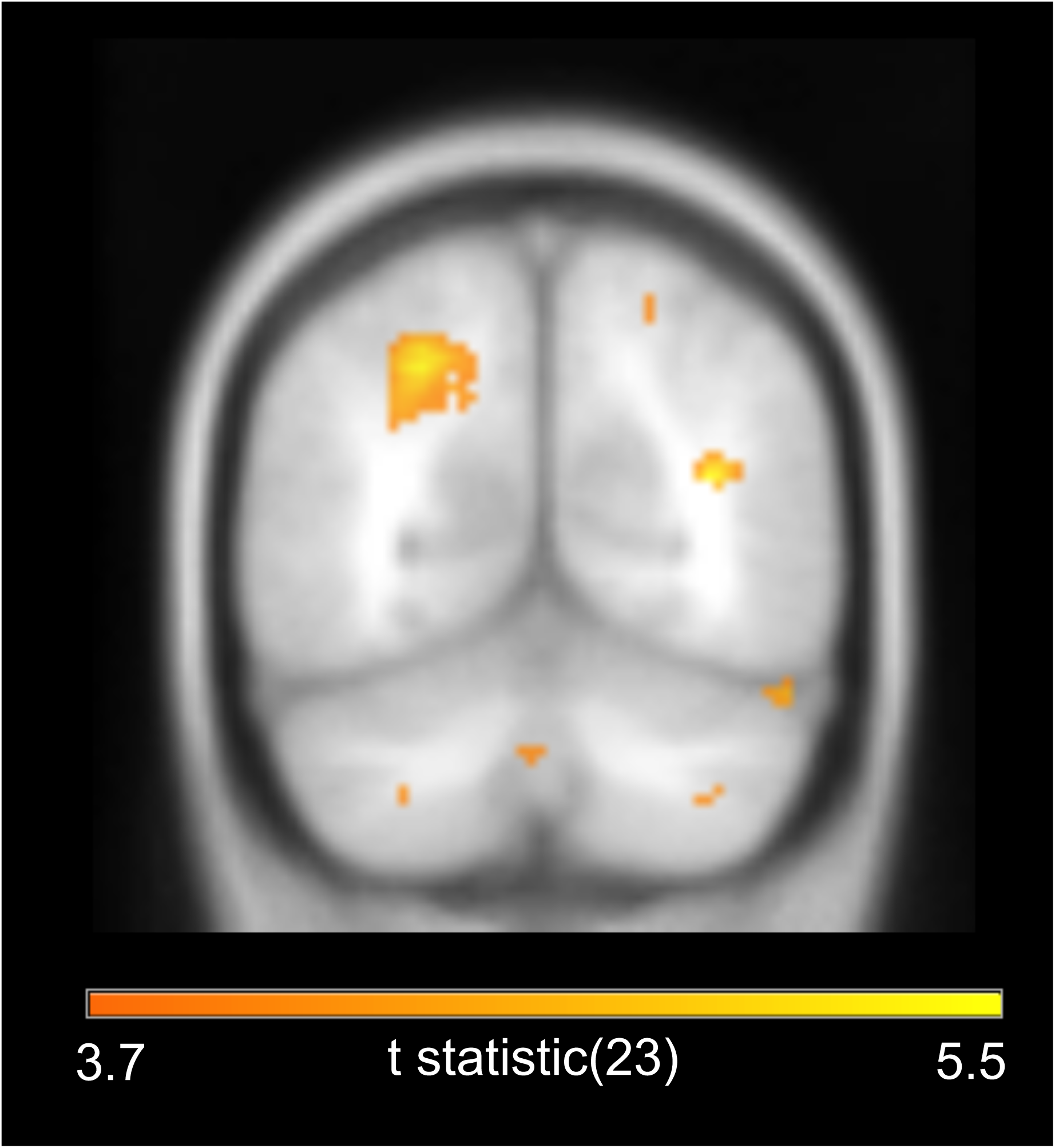
Coronal image of increased intraparietal sulcus activity for correct versus incorrect choices. Highlighted region survived cluster-level FWE correction at p< 0.05 and image is displayed at an uncorrected statistical threshold of p<.001.

**Figure 4-figure supplement 3.**
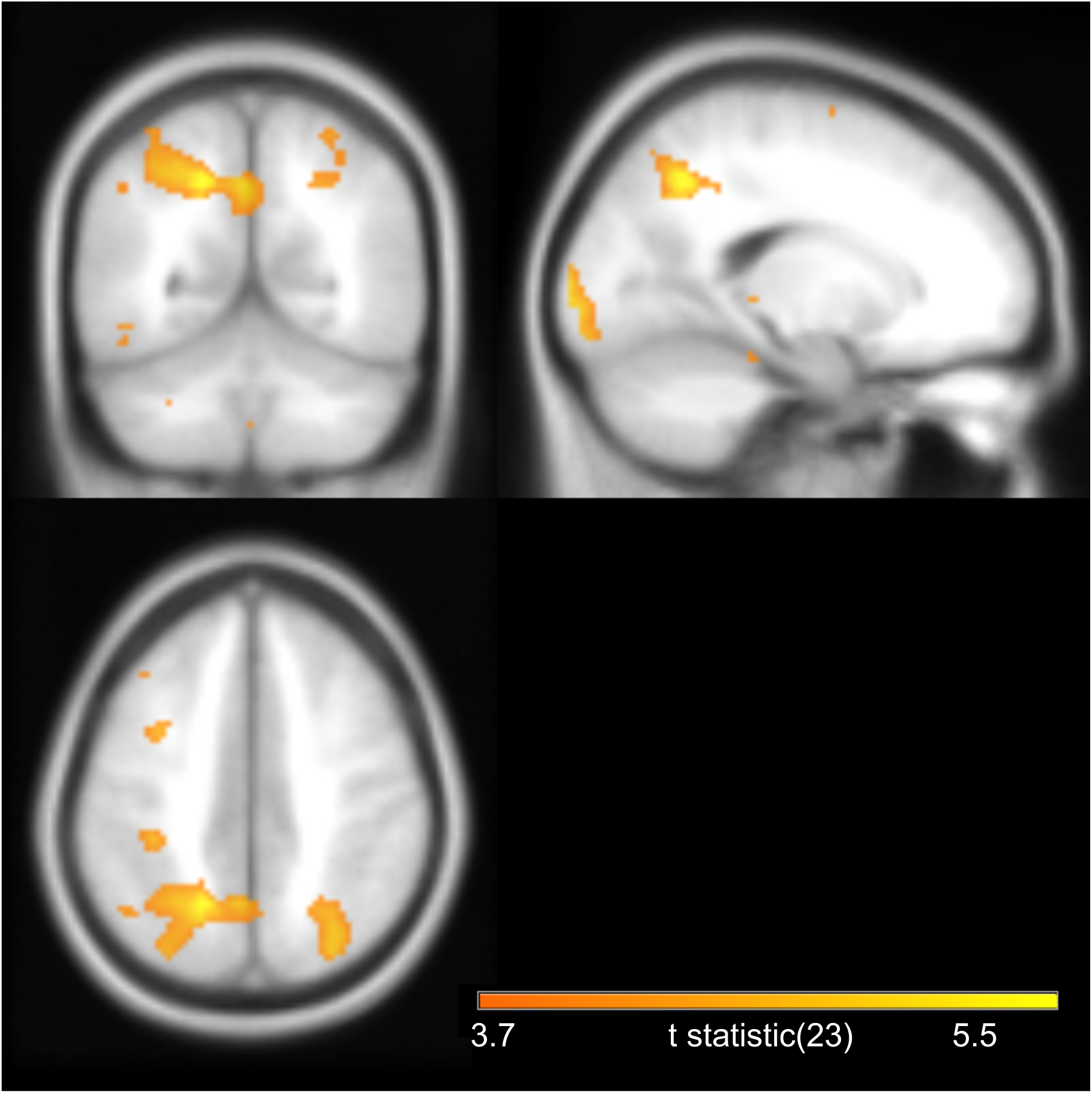
Images of increased posterior parietal cortex and precuneus activity for slower reaction times (RTs). Highlighted region survived cluster-level FWE correction at p< 0.05 and image is displayed at an uncorrected statistical threshold of p<.001.

## Source Data

Figure 1-Source Data 1. List of Rated Scenarios. Last 10 were used during practice trials and first 100 were used for the fMRI task.

Figure 2-Source Data 1. Data used to make Figure 2A.

Figure 2-Source Data 2. Data used to make Figure 2B.

Figure 2-Source Data 3. Data used to make Figure 2C.

Figure 2-Source Data 4. Data used to make Figure 2D.

Figure 2-Source Data 5. Data used to make Figure 2E.

Figure 2-Source Data 6. Data used to make Figure 2F.

Figure 2-Source Data 7. Data used to make Figure 2-Figure supplement 1.

Figure 2-Source Data 8. Data used to make Figure 2-Figure supplement 2.

